# Precise Control of Switchable Chimeric Antigen Receptor T Cells Allows Enhanced Safety and Less T Cell Exhaustion

**DOI:** 10.64898/2025.12.07.692875

**Authors:** Zihan Anna Zhang, Logan Herring, Thuzar Hla Shwe, Yue Hu, Xiaotong Song, Wenyue Cao, Wenshe Ray Liu

## Abstract

**Background:** Chimeric antigen receptor (CAR)-T cell therapies have achieved remarkable success in hematologic malignancies, yet their clinical utility remains limited by safety concerns, limited persistence, and T-cell exhaustion driven by continuous receptor signaling. While switchable CAR designs provide external control, many reported systems are irreversible, strictly binary, or compromise CAR-T potency.

**Methods:** We engineered an optimized chemically switchable CAR platform (CSN CAR) that pharmacologically regulates antigen engagement by controlling the surface expression of the full-length CAR. An NS3 protease module was embedded within the CAR construct to enable drug-dependent stabilization of intact CAR on T cells. Using engineered CAR-T cells, we quantified drug-controlled activation, cytotoxicity, and cytokine release against CD19^+^ tumor cells by flow cytometry and ELISA *in vitro*. We further screened clinically approved NS3/4A inhibitors in CAR-HEK and CAR-T cells to identify optimal small-molecule controllers. A chronic stimulation model was established to assess CAR-T persistence and exhaustion-associated phenotypes *in vitro*.

**Results:** CSN CAR-T cells enabled precise, dose-dependent regulation of CAR surface density, cytokine production, and cytotoxicity. In the OFF state, switchable CAR-T cells showed minimal basal activity, consistent with reduced antigen-driven activation and cytokine release in the absence of drug in the experimental conditions. Upon drug addition, intact surface CAR was detectable within 1 hour, reaching ∼80% of peak observed CAR expression by 4 hours. Reversible suppression of CAR expression enabled attenuation of cytotoxicity toward normal CD19^+^ B cells *in vitro* after target-cell reduction, supporting a potential strategy to mitigate prolonged on-target/off-tumor activity. Under chronic stimulation, switchable CAR-T cells exhibited reduced exhaustion-associated markers, more stable CAR expression, and preferential differentiation toward a central memory phenotype.

**Conclusion:** Together, these findings establish CSN CAR as a reversible and tunable switchable CAR-T platform enabled by clinically approved NS3/4A inhibitors, supporting controllable modulation of CAR activity with potential applications for improving the precision and safety of CAR-T cell therapy.

## Introduction

Chimeric antigen receptor (CAR)-T cell therapy is a type of immunotherapy that has emerged as a revolutionary approach for treating hematopoietic cancers, demonstrating remarkable clinical efficacy. CARs are genetic fusions of an antigen-recognizing extracellular fragment with a transmembrane (TM) and intracellular component for triggering T cell responses. T cells that are genetically engineered to express CARs recognize, bind, and eliminate cells expressing target antigens(1). The unprecedented success has led to the approval of seven CAR-T therapies by the U.S. Food and Drug Administration (FDA)(2-8). Despite these advances, CAR-T cell therapy faces several major challenges that need to be addressed. Severe and potentially life-threatening CAR-T cell-specific toxicities, such as cytokine release syndrome (CRS) and immune effector cell-associated neurotoxicity syndrome (ICANS), can occur. For currently approved CAR-T cell therapies, CRS incidence rate ranges from 42% to 95%(9). In addition, CAR-T cell efficacy is limited for specific tumor types, and tumor relapse remains a concern due to factors such as antigen escape and T cell exhaustion. In clinical data, anti-CD19 CAR-T cell therapy achieves outstanding response rates of 70% to 90% in pediatric B-cell acute lymphoblastic leukemia, but with high relapses(10-12). Continuous antigen stimulation contributes to CAR-T cell exhaustion, characterized by impaired proliferation and effector function(13). As such, enhancing CAR-T persistence is critical for improving long-term tumor control and reducing relapse.

A variety of current strategies could potentially address major limitations of CAR-T cell therapies. To diminish the impact of antigen escape, dual-CARs were designed(14-16). Alternatively, combination approaches, such as pairing CAR-T therapy with chemotherapy, can further enhance anti-tumor responses(17). To enhance CAR-T cell activity in an immunosuppressive environment, CAR design innovation is a favorable strategy(18, 19). For CAR-T cell-associated toxicity, it can be mitigated by altering the CAR structure to decrease the affinity of CAR receptors, modifying CAR spacers, replacing costimulatory domains, or selecting hyper-specific receptors for tumor antigens(20-24). To address CRS, inhibiting IL-6 secretion by macrophages and monocytes, or injecting IL-6 receptor antibodies, has been employed(25). To combat CAR-T cell exhaustion, PD-1 blockade can be applied(26). Additionally, more advanced designs of switchable CARs have been developed to limit excessive antigen engagement to delay T-cell exhaustion(27-34). However, many of the aforementioned methods have inherent limitations. For example, lowering CAR receptor binding affinity reduces potentially off-tumor toxicity, but it can also compromise tumor elimination. Similarly, inhibiting macrophage and monocyte activity may lower the risk of CRS, but it can also potentially impair innate immunity and delay tissue repair. The use of IL-6 receptor antibodies is effective in controlling CRS but may increase susceptibility to infections and obscure inflammatory symptoms(35). Among these strategies, switchable CAR-T cells are particularly promising because they allow post-manufacturing control over activity without broadly suppressing immune function, thereby mitigating selection pressure for antigen escape and avoiding T-cell overactivation.

In previously reported inducible CAR-T cells, distinct engineering strategies have been developed to achieve pharmacological control over CAR-T cell activity. For example, SNIP CAR(29) regulates CAR function through CAR intracellular signaling modulation by a multiplex, drug-gated circuit design. Similar with SNIP CAR, VIPER CAR(30) also achieves conditional T cells activation over intracellular signaling. The designs allowing programmable activation logic but requiring multi-component system architecture and maintaining CAR receptor presence at the cell surface regardless of control state. In comparison, our CSN CAR system adopts a fundamentally different strategy by directly regulating surface CAR receptor expression through protease-mediated cleavage. In this design, CAR receptors are removed from the cell surface in the absence of drug and stabilized only upon protease inhibitor treatment, thereby controlling antigen engagement at the receptor level rather than modulating downstream signaling. This streamlined, single-component architecture enables reversible and dose-dependent control of CAR availability. Importantly, by eliminating surface CAR expression in the OFF state, this approach may reduce unintended antigen engagement, minimize OFF-state leakage, and avoid potential tumor antigen masking observed in systems where CAR receptors remain present but inactive.

Nevertheless, switchable CAR-T cells have often been reported to exhibit reduced potency compared to conventional CAR-T cells (Conv.). We previously published an NS3/4A chemogenetic switch for CAR-T cells in which activity is regulated *in vivo*(36). Hepatitis C virus (HCV) NS3/4A protease is well characterized and has been extensively targeted in the development of antiviral drugs. Several small-molecule inhibitors have been approved for the treatment of HCV infection. However, this published chemogenetic CAR-T cells exhibited reduced CAR expression and diminished anti-tumor activity compared with Conv. CAR, which reduced overall therapeutic efficacy and compromised the translation potential. In this study, we designed a robust switchable CAR-T platform that enables control of T-cell activity without compromising tumor immunity and screened eight clinically approved NS3/4A inhibitors to identify effective and safer controller drugs.

## Methods

### Cell lines and cell culture

The cell lines HEK293T, K562, and Raji were obtained from American Type Culture Collection (ATCC). Primary cells Human Peripheral Blood Mononuclear Cells (PBMCs) were purchased from STEMCELL (70025.3) and activated by CD3/CD28 beads (ThermoFisher Scientific, 11131D) for further transduction. HEK293T cells were cultured in Dulbecco’s Modified Eagle’s Medium (DMEM, Corning, 10-013-CV) supplemented with 10% FBS (Gibco, 16000044). K562, Raji, and PBMC cells were incubated in RPMI 1640 (Corning, 10-040-CV) supplemented with 10% FBS (Gibco, 16000044). 100 U/ml IL2 was also supplemented in RPMI for T cell culture (PeproTech, 200-02).

### Generation of CAR constructs

The retroviral vector backbone was derived from Moloney murine leukemia virus and has been validated in clinical settings. It was generously provided by Dr. Andras Heczey of Baylor College of Medicine. The retroviral vector was designed to integrate CARs into primary human T cells, and it encodes our three sCAR designs (C&N, CSC, and CSN CAR) and one conventional CAR in this paper. The CAR gene fragments were synthesized by Twist Bioscience and were inserted into the retroviral vector in the lab. To generate switchable CARs, a CD19-targeting CAR receptor derived from an FDA-approved CAR (Kymriah), the sequence of NS3/4A and its cleavage site were adopted from a protein engineering paper(37). The conventional CAR has the same gene elements except for NS3/4A and its cleavage sites. The sequences of the CD8 hinge domain, the CD28 transmembrane domain, and the intracellular domains (CD28, 4-1BB, and CD3ζ) were from our lab and shown in this paper (Table S2). These three switchable CAR constructs contain the same intracellular signaling domains as the corresponding third-generation conventional CAR.

### Production of retrovirus and transduction of HEK293T and primary T cells

HEK293T cells were cultured three times before transient transfection with PEI-MAX (Polysciences, 49553-93-7), which transfects the retro-vector and CARs (C&N, CSC, CSN, or conventional CAR)- encoding plasmids into HEK cells. Virus-containing supernatant was collected after 48 and 72 h after transfection and filtered through a 0.45 μm filter. The supernatant was mixed with PEG-8000 at a 1:4 ratio and put at 4 degrees overnight. The virus was concentrated by centrifuging the PEG-8000-containing supernatant at 4000g for 2 h at 4 °C. After centrifuging, the supernatant was discarded, and the viruses were dissolved in PBS and stored at - 80 °C. To generate CAR-HEK cells and CAR-T cells, the HEK cells or CD3/CD28-activated PBMCs were incubated with concentrated virus and 8 μM polybrene, and then centrifuged with viruses at 1200g for 1 h, and then continued to culture the cells for 48 h. CAR-T cells were cultured in IL-2-containing media and maintained at 0.5-3*10^6 cells/ml. CAR-T cells are ready for analysis 7 days after transduction.

### Flow cytometry

The CAR expression level and positive percentage on CAR-HEK cells were detected by F(ab’)_2_ Fragment Rabbit Anti-Mouse IgG (H+L) (JacksonImmunoResearch, 315-006-003) antibody in flow cytometry. 5*10^4 CAR-HEK cells were stained with 2 μl antibody in 100 μl FACS buffer (PBS+4% FBS) at room temperature for 20 minutes and washed twice before flow cytometry analysis. The CAR expression level and positive percentage of CARs on CAR-T cells were identified by incubating 5*10^4 CAR-T cells with 2 μl PE-CD19 (ACRO Biosystems, CD9-HP2H5) protein in 100 μl FACS buffer at 4 °C for 1 h. The labelled CAR-T cells were washed twice before flow cytometry analysis. The antibodies used to characterize T cell phenotypes (table S1) were used to stain CAR-T cells according to the manufacturer’s instructions. The intracellular target detection on CAR-T cells, including antibody PE-human CD3, BV510-human IL-2(Biolegend, 500338), APC-human/mouse Granzyme B (Biolegend, 372203), and APC/Cyanine7-human Perforin (Biolegend, 308127). CAR-T cells stained with CD3 antibody, then washed with FACS buffer and fixed with fixation Buffer (Biolegend, 420801) for 20 minutes. After washing, cells were permeabilized and labelled with intracellular staining permeabilization wash buffer (Biolegend, 421002) according to the manufacturer’s instructions.

### Cytotoxicity assay

Cytotoxicity assays were conducted by incubating NT or CAR-T cells with Raji tumor cells at various effector-to-target (E:T) ratios (1:1, 5:1, 10:1) for 4 h, 18 h, or 48 h in an incubator with 5% CO_2_ at 37 °C. E:T ratio is based on CAR+ cells. Raji cells were replaced with K562 cells in a negative killing control group. At the beginning of incubation, 5 μM GZR was incubated to turn on the sCAR-T cells, and DMSO was added as a vehicle control. To evaluate dose-dependent T cell cytotoxicity, NT or CAR-T cells were incubated with Raji cells at a 1:1 E:T ratio for 48 h along with 0, 1, 10, 100, 1000, or 5000 nM GZR. After incubation, the total cell numbers were quantified by an auto-cell counter, and the cell mixture was labelled with PE-CD3 antibody to distinguish T cells and tumor cells. The percentage of tumor cells obtained from flow cytometry times the total cell number showed the residual tumor number. The supernatant was collected for the cytokine release assay.

### Cytokine release assay

To measure the level of released cytokine, including human TNF-α, IFN-γ, and IL-6, the supernatant was collected after the tumor killing assay. The ELISA kits ClinMax Human IL-6 ELISA Kit (ACRO Biosystems, CRS-B001), ELISA MAX-Human TNF-α (Biolegend, 430204), and ELISA MAX-Human IFN-γ (Biolegend, 430104) were used according to the manufacturer’s instructions. ELISAs were performed with 96-well Nunc MaxiSorp ELISA Plates (Biolegend, 423501).

### Primary macrophage and tumor cell triple-culture model

To better understand the role of sCAR-T cells in mediating the CRS-associated cytokine, IL-6. We did triple culture and detected macrophage-associated IL-6 to mimic CRS *in vitro*. Firstly, we separated monocytes from human PBMCs through adherence-mediated separation. We diluted PBMCs at 5×10^6 cells/ml in FBS-free RPMI-1640 and transferred cells to a cultured-treated plate. Put the plate at 37 degrees incubator for 2 h to get attached monocytes. Aspirate supernatant and wash the cells with warm PBS 3 times. Add 2 ml/well complete media (RPMI1640+10% FBS) and M-CSF at 50 ng/ml for macrophage differentiation. Replace half the media every 3 days and culture them for 7 days. After 7 days, the morphology of macrophages was confirmed using a Leica Dmi1 microscope equipped with a 40x objective lens. After macrophage differentiation, we incubated human primary macrophages with Raji tumors and CAR-T/NT cells at a 1:1:1 ratio for 24 h. After the coculture, the supernatant was collected, and the IL-6 levels were measured by ELISA.

### Drug-induced CAR expression on CSN CAR-T cells *in vivo*

CAR-T cells were generated and expanded *in vitro* as described above. After 8 days of culture, CSN CAR-T cells were intravenously infused into NSG mice at 5×10^6 CAR+ T cells per mouse (n=3). Five-week-old female NSG mice (Prkdc<scid>, Il2rg<tm1Wjl>) were purchased from The Jackson Laboratory. The animal experimental protocols were reviewed and approved by the Institutional Animal Care and Use Committee (IACUC) of Texas A&M University (Protocol No. 2025-0218 H). The NSG mice were housed strictly in a Specific Pathogen-Free (SPF) barrier facility at TAMU animal facility, fed with sterile water and an autoclaved (or irradiated) high-efficiency rodent diet. Peripheral blood samples were collected 2 days after infusion into EP tubes containing approximately 1 mL EDTA-DPBS (5 mM EDTA in DPBS) for baseline CAR expression analysis. Collected blood samples were centrifuged at 500 × g for 5 min, and the supernatant was removed. Cell pellets were resuspended in 1.2 mL ACK lysis buffer and incubated at room temperature for 15 min to lyse red blood cells. Following lysis, cells were centrifuged again and stained with anti-human CD3 antibody and PE-conjugated CD19 protein at 4°C in the dark for 1 h. Cells were washed twice with FACS buffer before flow cytometry analysis. For drug administration, GZR was initially dissolved in DMSO as a stock solution and subsequently formulated in a vehicle containing 5% DMSO, 40% PEG300(MCE, HY-Y0873), 5% Tween-80(MCE, HY-Y1891), and 50% DPBS(Corning, 21-031-CM). Mice received GZR treatment at 50 mg/kg by daily intraperitoneal injection for 2 consecutive days, followed by repeat peripheral blood collection and CAR expression analysis by flow cytometry.

### Chronic stimulation model

*In vitro* expansion and persistence of CAR-T cells were evaluated by a chronic stimulation model. NT or CAR-T cells were cocultured with fresh Raji tumor cells for 3 rounds, each round allows a 48 h killing. The fresh tumor cells were added at every round at E:T 1:1 ratio. The 2nd round started right after the 1st round, and there is a four-day rest between the 2nd and 3rd rounds. T cell cytotoxicity was evaluated after each round, and T cell activation, differentiation, exhaustion, and positive CAR expression were assessed after the 3rd round.

### Statistical Analysis

Two experimental groups were compared using an unpaired two-tailed t-test. *P*-values were calculated with GraphPad Prism 10.6.1 and statistics represented by not significant (ns) = *p* > 0.05, * = *p* < 0.05, ** = *p* < 0.01, *** = *p* < 0.001. All error bars are represented either SEM or SD.

## Results

### Design and optimization of reversibly switchable CARs

We aimed to develop a switchable CAR (sCAR)-T cell system whose activation can be reversibly controlled by regulating surface CAR expression. Additionally, we intended to equip CAR-T cells with tunable cytokine release and anti-tumor immunity. To achieve this, we genetically fused an NS3/4A protease complex into a Conv. CAR as a molecular switch. NS3 cleavage sites (CSs) were also introduced into the CAR to enable NS3/4A to cleave CAR receptors from the cell surface. Inhibition of NS3 activities stabilizes intact CARs, allowing them to bind antigen and initiate downstream signaling. Thus, sCAR-T cells are in an OFF state by default and can be switched ON by applying NS3/4A inhibitors. sCAR expression and function are therefore expected to be regulated by small-molecule inhibitors (Figure 1A). To build the sCAR construct, we used an anti-CD19 CAR receptor derived from Kymriah, together with the TM domain, followed by intracellular costimulatory domains including CD28 and 4-1BB, and a CD3ζ signaling module. In our sCAR designs, the NS3/4A switch and its CSs were incorporated into the extracellular region of the CAR construct to maintain normal intracellular signaling architecture while enabling regulation of surface CAR presence. To systematically assess cleavage efficiency, we generated three CAR variants: (i) CSs at both the *N*- and *C*-termini of switch (C&N CAR), (ii) a single *C*-terminal site (CSC CAR), and (iii) a single *N*-terminal site (CSN CAR) (Figures 1B–D). All were designed to remain in the OFF state without an inhibitor and differ in cleavage position, which may influence sCAR stability and regulatory efficiency.

**Figure 1.**
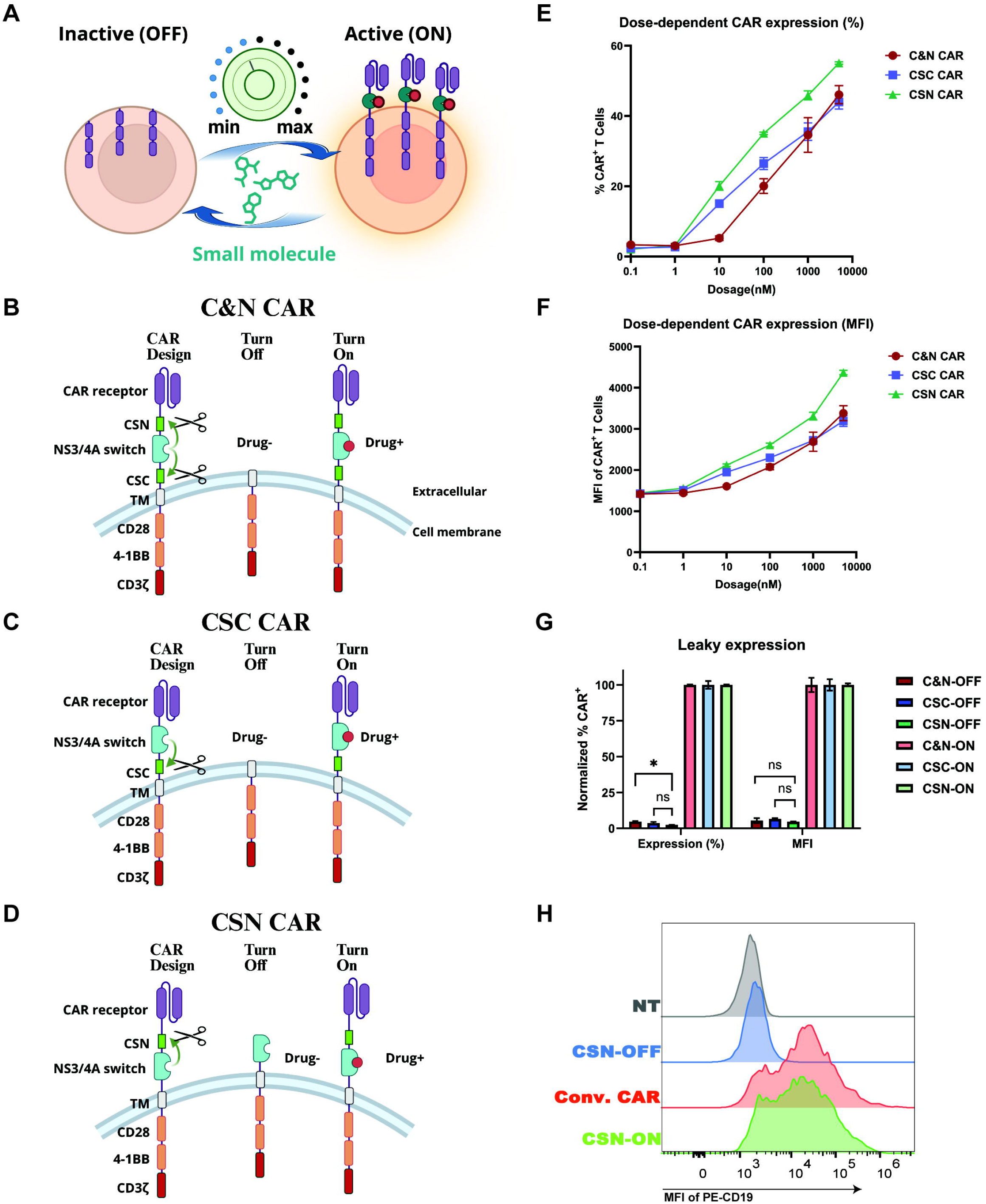
Designs of switchable CAR (sCAR) T cells and characterizations of their sCAR control ranges. (**A**) Schematic diagram illustrating the design of a precisely controlled CAR-T cell system. The sCAR-T cell is inactive (OFF state) in the absence of a small molecule drug inhibiting a switch integrated in CAR. It can be turned ON by adding a small molecule drug, allowing dynamic control. (**B**) Design of C&N CAR and its controlled CAR expressions in OFF and ON states. NS3/4A, as a switch, cleaves at both *N*- and *C*-termini in the OFF state. (**C**) The design and expression format of CSC CAR on T cells in the OFF and ON states. The NS3/4A switch only cleaves at its C-terminal in the OFF state. Conv. CAR-T cells continuously express full-length CARs and have the same components as sCAR except for the NS3/4A switch and CSs. (**D**) The design and expression format of CSN CAR on T cells in the OFF and ON states. NS3/4A switch only cleaves at its N-terminal in the OFF state. B, C, and D were generated by Biorender. (**E**) The comparison of CAR expression percentage between C&N, CSC, and CSN CAR designs. Human CAR-T cells were treated with DMSO or various doses of small-molecule drugs for 24 hours. Phycoerythrin (PE)-conjugated CD19 protein (PE-CD19) was used to label functional binding CAR-T cells. CAR expression levels were characterized by median fluorescence intensity (MFI) of PE-CD19. The positive CARs were detected by flow cytometry (mean ± SEM, n = 3). (**F**) The comparison of CAR expression level (MFI) between C&N, CSC, and CSN CAR designs for 24 h. The human CAR-T cells were cultured in DMSO or various concentrations of small-molecular-weight drugs (mean ± SEM, n = 3). (**G**) The leakage of sCAR expression in the OFF state was identified. C&N, CSC, and CSN CAR-T cells were incubated with DMSO for the OFF group or 5 μM GZR for the ON group for 24 h. The expression of percentage and MFI was normalized to the ON group (mean ± SEM, n = 3). (**H**) The comparison of the maximum CAR expression level of CSN-ON CAR-T with Conv. CAR, and a minimum of CSN CAR with NT. Non-transduced (NT) T cells and Conv. were incubated with DMSO. The expression level (histogram) was displayed with a staggered offset. Part of the figure was generated by Biorender.

To evaluate regulatory performance, we cloned the three constructs and generated sCAR-T cells separately. sCAR-T cells were then cultured with a clinically approved NS3/4A inhibitor (grazoprevir, GZR). After drug treatment, we assessed the percentage and level of CAR expression by incubating CAR-T cells with PE labelled CD19 protein (PE-CD19) and then analyzing them using flow cytometry. The positive percentage and median fluorescence intensity (MFI) of PE on T cells represent positive CAR percentage and expression level. All three designs of sCAR-T cells showed dose-dependent CAR expression percentage and level (Figures 1E and 1F). In comparison, CSN CAR displayed the most sensitive response to the controller drug, as indicated by both positive CAR percentage and expression level (MFI). To better understand potential leaky activity due to residual CAR expression, we cultured the three types of sCAR-T cells without a controller drug to turn their switches OFF and compared them with non-transduced T cells (NT). The three designs showed no apparent leakage, and CSN and CSC had lower CAR+ percentages in their OFF state, indicating tighter suppression of CAR surface expression in the absence of drug. (Figures 1G and S1). Importantly, the CSN CAR demonstrated the strongest inducible CAR expression in response to the drug. Therefore, the CSN CAR provided an optimal balance between minimal leakage (Figure S1) with a slight MFI change relative to NT (Figure S2) and robust inducible CAR expression. To determine whether infusing the switch impairs CAR expression, we next turned ON the CSN CAR-T cells (CSN-ON) and compared their CAR expression level (MFI) with Conv. CAR-T cells. CSN-ON exhibited a comparable CAR expression level to the Conv. CAR, and CSN CAR-T cells at the OFF state (CSN-OFF) showed a similar CAR expression level to NT (Figure 1H). To evaluate whether incorporation of the NS3/4A switch intrinsically affects CAR-T cell expansion, we compared the cell growth rate of CSN and Conv. CAR T cells under *in vitro* culture conditions. No significant differences in absolute T-cell expansion were observed between these two groups (Figure S3), suggesting that the NS3/4A switch does not measurably alter basal CAR-T cell growth. Overall, CSN CAR demonstrated the widest range of control and is optimal compared to the other two, enabling controllable, dose-dependent full-length CAR expression without compromising their levels. As such, we moved forward with this CSN CAR design for subsequent experiments.

### Drug screening of NS3/4A on CSN CAR-expressing cells for more efficient control

NS3/4A is a serine protease essential for HCV polyprotein processing. Multiple generations of inhibitors have been developed for NS3/4A, progressing from early peptidomimetics to more advanced macrocyclic compounds with improved potency and stability. The U.S. FDA and other regulatory agencies have approved several inhibitors. The availability of diverse, clinically validated small-molecule inhibitors makes NS3/4A an attractive module for synthetic biology applications, such as CAR-T cells regulation. The inhibitors include first-generation inhibitors, linear peptidomimetic compounds, and second-generation macrocyclic inhibitors, such as ASV (asunaprevir). Additionally, the third-generation inhibitors, such as GZR (grazoprevir) and GLE (glecaprevir), feature optimized macrocyclic scaffolds.

To identify safe and effective small-molecule drugs for controlling the designed sCAR-T system, we manufactured HEK293T cells expressing CAR constructs (CAR-HEK) and initially screened clinically approved NS3/4A inhibitors on CAR-HEK cells, which provides a more cost-effective platform for characterizing the switchable system. In addition, tipranavir (an HIV-1 protease inhibitor), which shares similar molecular weight and hydrophobicity with NS3/4A inhibitors, was included as a negative control to test the specificity of CAR expression control. The CAR receptor domain is structured as a scFv (single-chain variable fragment), and we detected scFv levels on the HEK cell to evaluate the induced CAR expression level. In the absence of inhibitors, CAR-HEK cells failed to express CAR receptors. By contrast, incubation with NS3/4A inhibitors induced full-length CAR expression in a dose-dependent manner, whereas tipranavir didn’t trigger expression at any concentrations (Figure 2A). This result confirmed that full-length CAR expression is specifically regulated by NS3/4A inhibitors rather than by unrelated small molecules. Among the inhibitors tested, GZR, GLE, and ASV (the top three drugs) demonstrated the highest potency in promoting CAR expression. GZR and GLE induced a detectable full-length CAR expression at approximately 1 nM and achieved robust activity as low as 100 nM. ASV was less potent than GZR and GLE. The remaining inhibitors displayed lower potency but still produced measurable expression. To assess drug toxicity, HEK293T cells were treated with the nine inhibitors, and the top three drugs showed minimal effects on cell viability (Figure S4). Overall, these results demonstrated that NS3/4A inhibitors can efficiently regulate full-length CAR expression in CAR-HEK cells with low toxicity, validating the feasibility of the sCAR system. Given the well-regulated control of full-length CAR expression by the top three drugs, we next evaluated the time-dependent response of CAR expression to them (Figure 2B). Across the tested range, full-length CAR expression increased with both drug concentration and incubation time. Drug treatment induced detectable CAR expression within 1 hour, robust expression by 4 hours, and high CAR expression levels by 24 hours. GZR and GLE consistently induced higher CAR expression, highlighting their superior efficacy in regulating the sCAR system.

**Figure 2.**
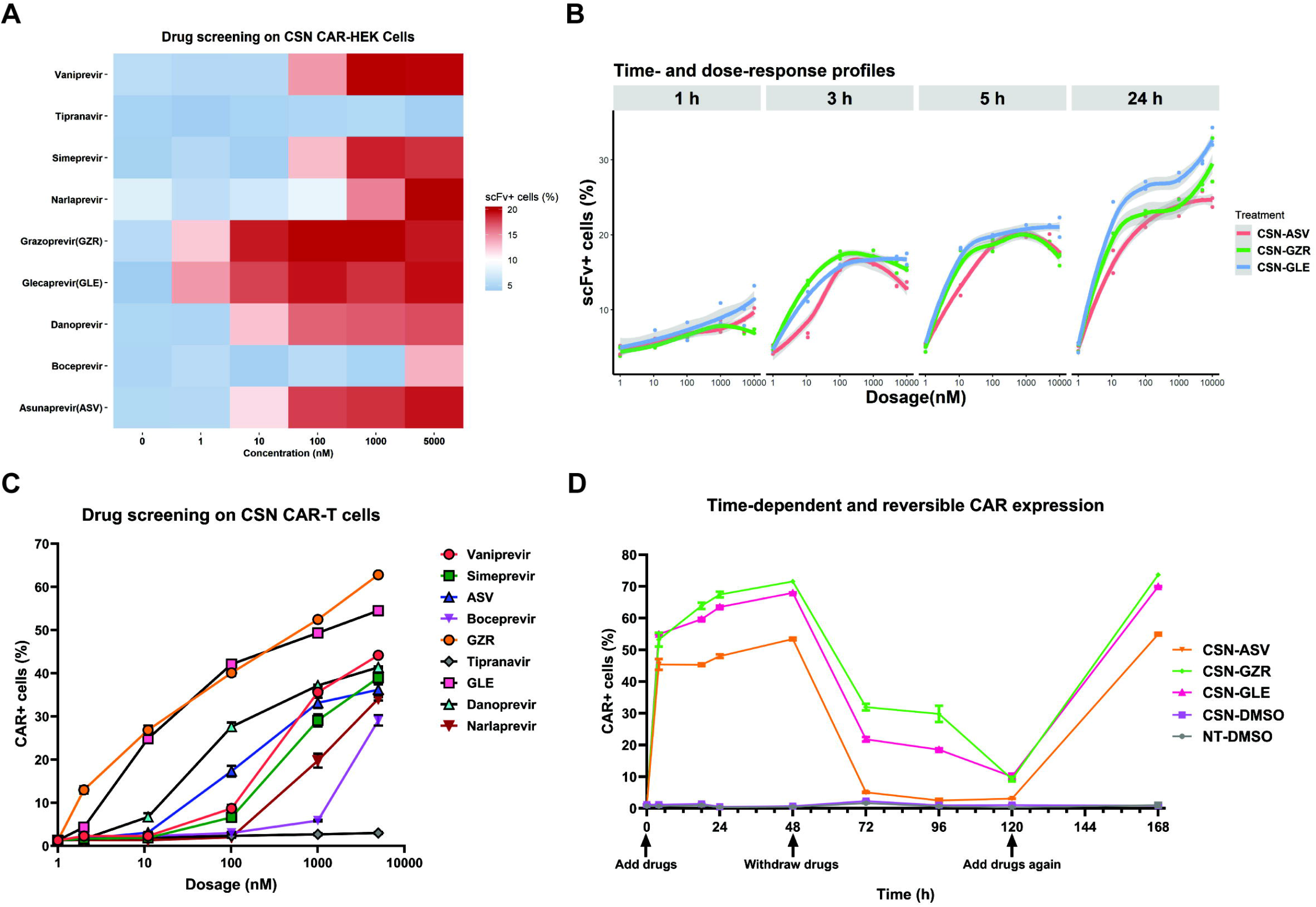
Drug screening of NS3/4A inhibitors on CSN CAR-expressing HEK293T cells (CSN CAR-HEK) and T cells (CSN CAR-T). (**A**) Screening of potent HCV NS3/4A inhibitors on CNS CAR-HEK cells. CAR-HEK cells are sCAR-expressing HEK293T cells. Drug-induced full-length CAR expression was compared by identifying positive full-length CAR in CNS CAR-HEK cells. CSN CAR-T cells were incubated with various inhibitors for 24 h. After incubation, the cells were trypsinized to detach from their culture dishes and collected. Cells were analyzed by flow cytometry through labelling with Alexa647-conjugated anti-scFv (structure of CAR receptor). The figure was plotted by ggplot2 by defining the highest detected level as dark red and scaling from 0 to 20. (**B**) The time and dose-response profiles of full-length expression responding to ASV, GZR, and GLE on CSN CAR-HEK cells. The clinic C_max_ of ASV, GZR, and GLE are ∼572, ∼165, and ∼597 ng/mL, respectively. The CAR-HEK cells were incubated with different inhibitor concentrations for 1, 3, 5, and 24 h separately, and then the full-length CAR expression levels were quantified. CNS CAR-HEK cells were stained with Alexa647-anti-scFv and analyzed by flow cytometry. The figure was plotted by ggplot2. (**C**) The drug screening of clinically approved HCV NS3/4A drugs on CSN CAR-T cells. The potency of drugs was evaluated by drug-induced full-length CAR receptor expression (mean ± SEM, n = 3). (**D**) The time-dependency of drug-regulated full-length CAR expression and reversible control of CSN CAR-T cells. The CSN CAR-T cells were incubated with 5 μM GZR for a time course to reveal time dependency. After 48 h of incubation, cells were washed three times with RPMI 1640 medium containing 10% FBS and cultured without drugs for an additional 48 h (mean ± SEM, n = 3).

After confirming the functionality of NS3/4A inhibitors in CAR-HEK cells, we next manufactured CSN CAR-T cells and evaluated the potency of inhibitors in human CAR-T cells (Figure 2C). Among the inhibitors, GZR and GLE again showed the highest potency in regulating CAR expression at all tested concentrations. The remaining inhibitors induced detectable but weaker responses, whereas tipranavir failed to trigger expression consistent with CAR-HEK cell-derived data. Additionally, we assessed the cytotoxicity of the top three drugs in T-cells and observed low toxicity (Figure S5). To further evaluate whether systemic inhibitor administration could induce the CSN platform *in vivo*, NSG mice infused with CSN CAR-T cells were treated daily with GZR (50 mg/kg). CAR surface expression on human T cells was analyzed before and after drug administration. GZR treatment induced a clear increase in CAR-positive T cells compared with untreated controls, demonstrating that systemic administration of the inhibitor can pharmacologically induce CAR expression *in vivo* (Figure S6).

To evaluate time-dependent CAR expression, CSN CAR-T cells were treated with drugs at various time points (Figure 2D). Full-length CARs were expressed within 4 h, reaching high CAR expression levels overnight, and continued to rise modestly over 24 h. To further assess the reversibility of CAR expression, drugs were washed out after certain drug incubation times, and cells were continued to be cultured without drugs. Withdrawing drugs led to a sharp decline in full-length CAR expression within 24 h. Reintroduction of drugs restored CAR receptor expression to similar levels before drug withdrawal, confirming that our designed sCAR system is both controllable and reversible (Figure S7).

### Drug-controlled cytotoxicity, cytokine release, and effector molecule expression in CSN CAR-T cells

To investigate whether CSN CAR-T cells regulate cytotoxicity, we first quantified residual tumor cells after coculturing with CAR-T cells at various E:T (Effector to Target) ratios. NT cells were included as negative controls for effector cells (Figures 3A-3C). After coculturing, Conv. CAR-T cells drove tumor clearance at all ratios and time points, consistent with their effective tumor suppression. After 18 h incubation or longer, CSN-ON CAR-T cells significantly eliminated Raji cells compared with CSN-OFF at all ratios, but not in 4 h, likely reflecting the time required for CAR expression. Strikingly, CSN-ON CAR-T cells reached or even exceeded the cytotoxicity of Conv. CAR at 18 h with a 10:1 ratio. In contrast, CSN-OFF CAR exhibited no killing, indicating minimal background leakiness. Together, these results suggest that tumor cytotoxicity of CAR-T cells is tightly controlled by CSN CARs at both high and low E:T ratios. We next explored the control of cytokine release, focusing on TNF-α, IFN-γ, and IL-6. These cytokines play crucial roles in anti-tumor activity but can also contribute to CAR-T cell-associated toxicity. Thus, monitoring and controlling these cytokines are key to balancing efficacy versus safety in CAR-T therapy. As described above, Raji cells were cocultured with CAR-T cells (Figures 3D-3F), the supernatant was collected after coculture to measure cytokine levels. CAR-T cells produced substantial levels of TNF-α at E:T ratios of 1:1 and 5:1 but much less at a high E:T ratio (10:1), likely because TNF-α release is rapid and transient compared to IFN-γ or IL-6, and target cells were cleared quickly at high effector density. CSN CAR-T cells showed controlled TNF-α levels, with apparent ON/OFF differences. For IFN-γ, both Conv. and CSN-ON CAR groups achieved higher levels than NT and CSN-OFF groups, indicating a controllable IFN-γ release in CSN CAR-T cells. Moreover, CSN CAR-T cells showed a well-controlled IL-6 cytokine release profile across all E:T ratios, suggesting a reduced risk of CRS-like toxicity. Overall, these results demonstrated that the key cytokines in CAR-T cell therapy are controllable in CSN CAR-T cells, suggesting a low risk of immune toxicity in a tumor-surrounding environment. IL-2 is an essential cytokine that supports T cell expansion and is supplemented in T cell culture media. Supernatant IL-2 does not specifically reflect cytokine release by CAR-T cells. To more accurately assess, we therefore analyzed the expression of IL-2, granzyme B, and perforin directly in CAR-T cells. After incubation, cells were stained with antibodies, fixed, and permeabilized to detect intracellular expression. The expression levels of IL-2, granzyme B, and perforin were assessed intracellularly (Figure S8), CSN CAR-T cells also exhibited well-regulated expression of them during tumor killing.

**Figure 3.**
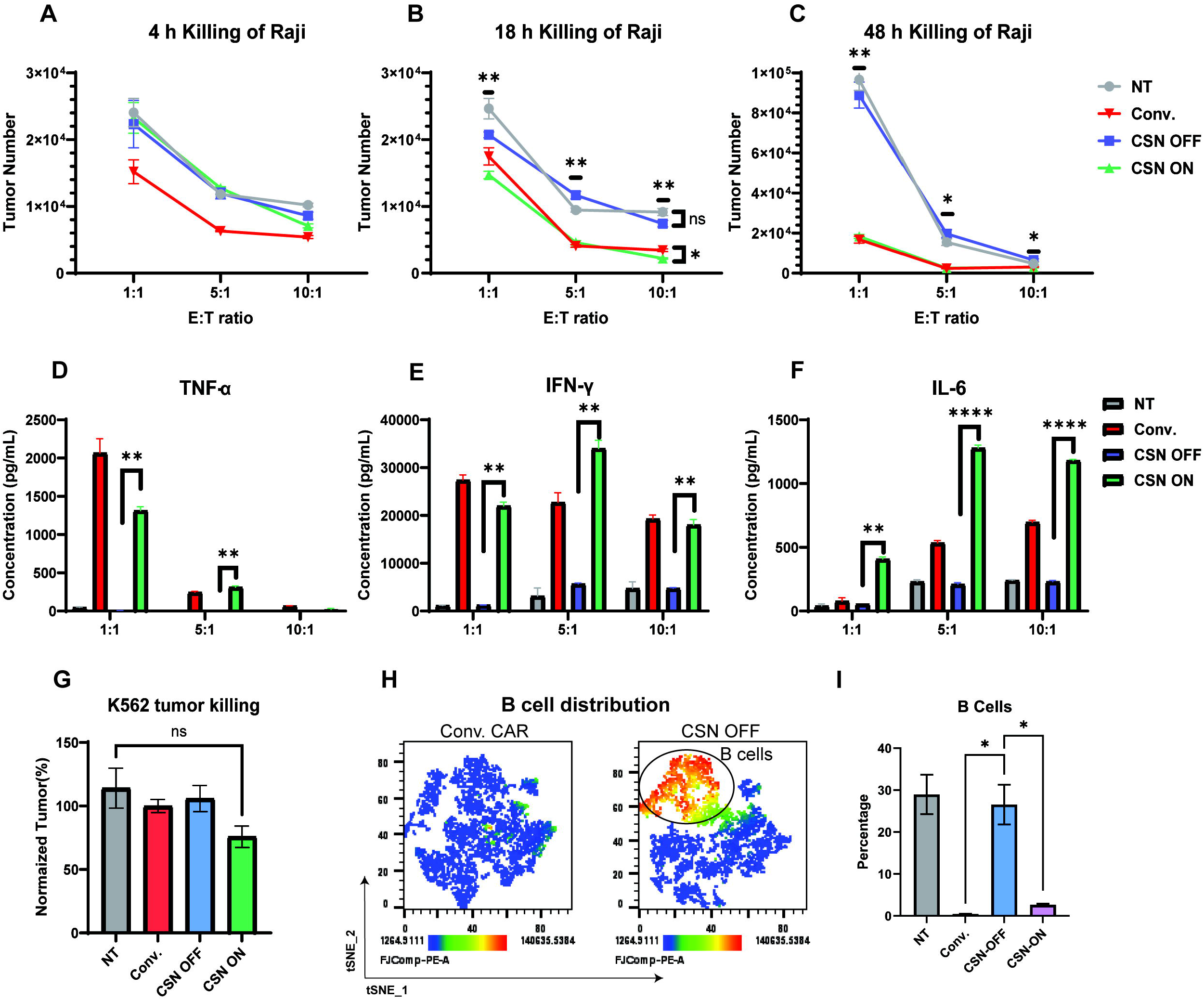
Characterization of the control of cytokine release, cytotoxic effector molecule expression, and cytotoxicity on CSN CAR-T cells. (**A-C**) The controlled cytotoxicity of CSN CAR-T cells. The Raji cells were cocultured with Conv. or CSN CAR-T cells and treated with DMSO or 5 μM GZR. For the CSN groups, cells were incubated with Raji cells and DMSO (CSN-OFF) or GZR (CSN-ON). Conv. CAR-T and NT cells were cocultured with Raji cells and DMSO as positive and negative controls, respectively. Effector-to-target (E:T) ratios of 1:1, 5:1, and 10:1 were tested for 4, 18, and 48 h (E:T ratio is based on CAR+ cells). After coculturing, the cell mixture was washed once with FACS buffer and labelled with CD3 antibody to distinguish tumor cells from T cells. The viability dye, HelixBlue, was added before running the flow cytometry without washing (mean ± SEM, n = 3). (**D-F**) The regulated cytokine release of CSN CAR-T cells. After coculturing Raji and CAR-T cells and drug treatment, the cell suspension was centrifuged to collect the supernatant, and cytokine levels were measured by ELISA (Enzyme-Linked Immunosorbent Assay). Tumor necrosis factor-alpha (TNF-α), interferon gamma (IFN-γ), and interleukin 6 (IL-6) (mean ± SEM, n = 3). (**G**) Killing of non-targeting K562 tumor cells (CD19^-^). K562 tumor cells were incubated with NT/CAR-T cells for 48 h. Tumor cell numbers were quantified after incubation and normalized to the control group. (**H**) B cell distribution in PBMCs after incubation with CAR-T cells. PBMCs and CAR-T cells were cocultured for 24 h. After incubation, the cell mixture was analyzed by flow cytometry, and the cells were labeled with CD3 and CD19 antibodies, B cells (CD3^−^CD19^+^). t-SNE shows the distribution of B based on the CD3^-^ cell populations. The color axis represents the expression level of CD19^+^ cells. B cells labelled in the figure are CD3^-^CD19^+^. t-SNE-1 and t-SNE-2 represent the first and second dimensions of the t-SNE embedding and do not correspond to defined biological variables. (**I**) The percentage of B cells (CD3^-^CD19^+^) in CD3^-^ cell populations after incubating PBMCs with NT or CAR-T cells. The total cells were labelled with both CD3 and CD19 antibodies and analyzed by flow cytometry.

To evaluate non-specific cytotoxicity from CAR-T cells, K562 cells (CD19^−^) were cocultured with CAR-T cells, K562 cells are also incubated GZR in the absence of T cells as a control group (Figure S9). Neither Conv. nor CSN CAR-T cells exhibited significant killing (Figure 3G). As expected, CD19-CAR-T cells efficiently eliminated CD19^+^ cells, however, CD19 is also critical and universally expressed on normal B cells. Hence, CD19-targeting treatment inevitably leads to depletion of healthy B cells even after tumor clearance and brings a variety of side effects. We next asked whether the OFF state of the CSN CAR would abolish on-target, off-tumor activity against normal CD19^+^ B cells after tumor clearance. To model a situation in which CAR-T cell activity is intentionally attenuated after disease control, we cocultured human PBMCs with CSN-OFF or CN-ON CAR-T cells and quantified residual CD19^+^ cells to detect the presence of B cells (Figures 3H and 3I). In this model, the CSN-OFF system effectively spared CD19^+^ B cells, whereas Conv. CAR-T cells induced near-complete B-cell depletion. These results demonstrate that pharmacological suppression of CAR expression can abrogate off-tumor cytotoxicity against healthy B cells following tumor clearance. Preserving the non-malignant B-cell population in this manner represents a viable strategy to mitigate treatment-induced B-cell aplasia. To further determine whether the OFF-state CSN CAR system masks tumor-associated CD19 epitopes, Raji tumor cells were cocultured with CSN-OFF CAR-T cells or NT cells, followed by staining with a flow cytometry anti-CD19 antibody whose recognition epitope substantially overlaps with the CAR scFv binding region (Figure S10). No detectable reduction in CD19 antibody binding was observed in the CSN-OFF group compared with NT controls, indicating that OFF-state CSN CAR-T cells do not measurably block or mask CD19 epitope accessibility on tumor cells. These findings are consistent with minimal residual antigen engagement in the OFF state.

### CSN CAR-T cells achieved precise control of CAR-T cell activation, tumor cytotoxicity, and cytokine release

After establishing that tumor cytotoxicity and cytokine release are controllable in the ON and OFF states of CSN CAR-T cells, we next investigated whether this regulation could also be graded. To evaluate modulation accuracy, CAR-T cells were cocultured with Raji cells at various GZR concentrations, and T cell activation, cytotoxicity, and cytokine release were then characterized. In the activation analysis, CSN-OFF CAR-T cells exhibited levels similar to NT, confirming that they could not be activated by surrounding tumor cells in their OFF state (Figures 4A and 4B). In parallel, both the percentage and level of activation showed dose-dependent increase with the provided GZR concentration. CSN CAR-T cells reached the activation level of Conv. CAR at around 1 μM GZR and exceeded it at higher concentrations. These findings demonstrated that the activation level is finely tunable by drug concentration under tumor antigen stimulation. We next examined whether tumor cytotoxicity can be dose-dependently controlled in our switchable system (Figure 4C). CSN-OFF showed minimal cytotoxicity similar to NT, confirming the absence of CAR-driven killing in the OFF state. With increasing GZR concentrations, cytotoxicity rose in a dose-dependent manner, eventually reaching Conv. CAR levels. This result underscores the switchable system’s ability to delicately regulate tumor killing by CAR-T cells.

**Figure 4.**
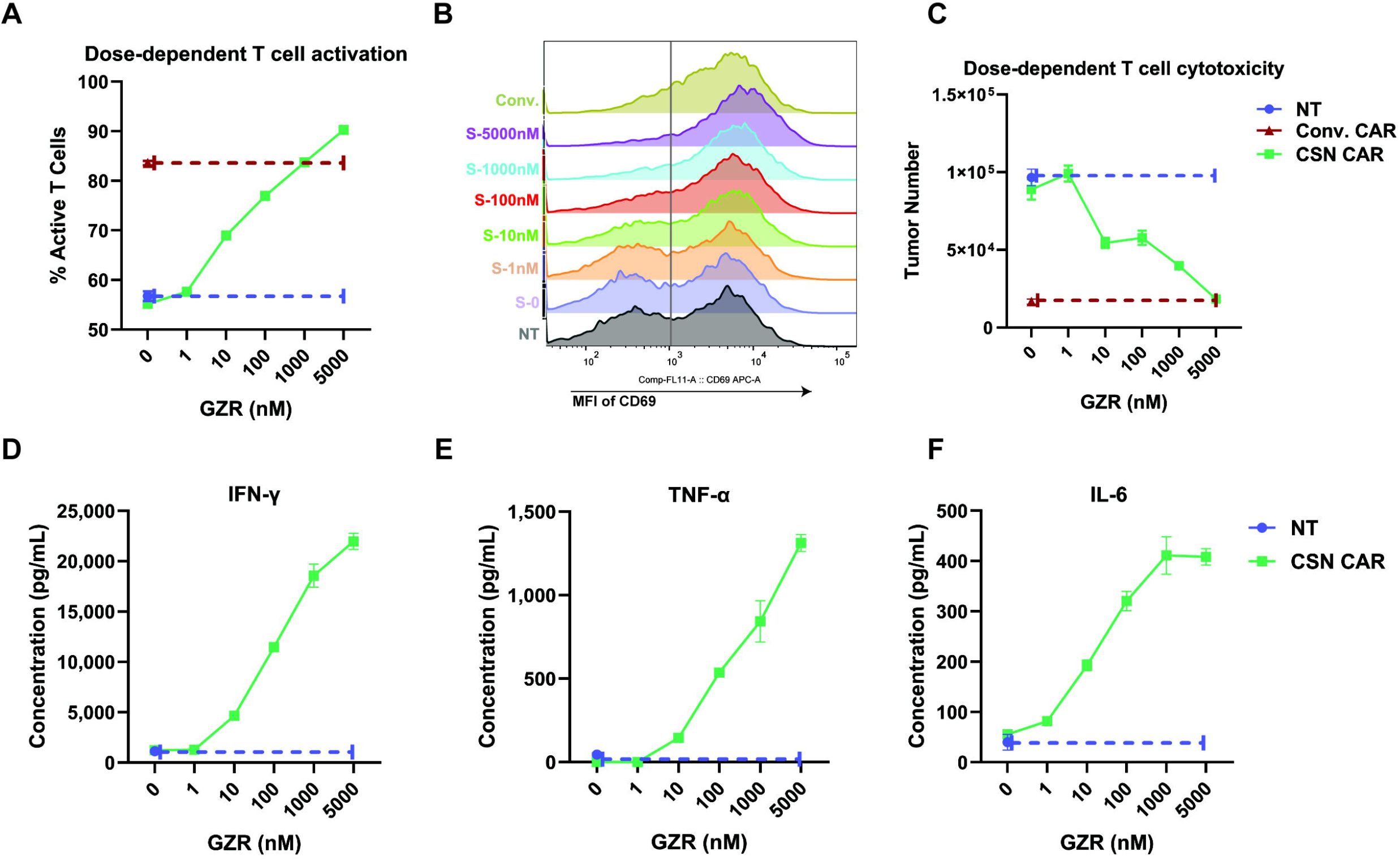
Identification of dose-dependent control of T cell activation, tumor cytotoxicity, and cytokine release in CSN CAR-T cells. (**A**) Percentage of activated CAR-T cells (CD3^+^CD69^+^) and (**B**) activation level (CD69 MFI). CD69 is a canonical activation marker. CSN CAR-T cells were cocultured with Raji tumor cells and either DMSO (OFF state) or increasing concentrations of GZR (1 nM–5 μM) for 48 h. Conventional (Conv.) CAR-T cells and non-transduced (NT) T cells were cocultured with Raji cells and DMSO for 48 h as positive and negative controls, respectively. MFI indicates the expression level of CD69 on CD3^+^ T cells. GZR concentrations: S-0 (DMSO only), S-1 (1 nM), S-10 (10 nM), S-100 (100 nM), S-1000 (1 μM), and S-5000 (5 μM). (**C**) Dose-dependent cytotoxicity of CSN CAR-T cells. Remaining tumor cell numbers were quantified after 48 h of coculture with CSN CAR-T cells and graded GZR concentrations. Conv. and NT-T cells cocultured with Raji and DMSO served as positive and negative controls. (**D–F**) Dose-dependent cytokine release (IFN-γ, TNF-α, and IL-6) by CSN CAR-T cells. Cells were cocultured with Raji cells and either DMSO or graded GZR concentrations for 48 h. Cytokine levels were quantified using ELISA kits.

Cytokines act as a double-edged sword in adoptive cell therapy, which are essential for antitumor activity, and their overproduction drives systemic toxicities such as CRS. Consequently, achieving a therapeutic window requires a delicate balance between efficacy and safety. To address this, we characterized cytokine secretion profiles of CSN CAR-T cells, focusing on IFN-γ, TNF-α, and IL-6, which are key mediators in CRS. In the absence of the drug, CSN-OFF CAR-T cells exhibited negligible cytokine release, confirming minimal basal leakage of the switch. Upon drug treatment, cytokine secretion increased in a dose-dependent manner. IFN-γ and TNF-α reached maximal levels at 5 μM GZR, while IL-6 peaked earlier at 1 μM and plateaued at higher concentrations (Figures 4D-4F). Because clinical cytokine release syndrome (CRS) is driven predominantly by activated myeloid cells rather than CAR-T cells directly, we next evaluated whether pharmacologic tuning of CSN CAR-T activity modulates secondary macrophage-associated IL-6 secretion. To model this process, we separated monocytes from PBMCs and differentiated them into macrophages by adding M-CSF for 7 days. After macrophage differentiation, primary human macrophages were cocultured with Raji tumor cells and CAR-T cells at varying GZR concentrations (1–100 nM). Supernatants were collected, and IL-6 levels were quantified by ELISA (Figure S11). Compared with Conv. CAR-T cells, CSN CAR-T cells in the OFF or low-dose ON states induced substantially lower IL-6 secretion from macrophage-containing cocultures. These findings indicate that pharmacologic attenuation of CSN CAR-T activity can reduce secondary inflammatory signal (IL-6) from macrophages *in vitro*. Unlike the constitutive and uncontrollable activity of conventional CAR-T cells, our switchable platform enables tunable and predictable regulation. These results demonstrate that CSN CAR-T cells provide precise, graded control over T-cell activation, cytotoxicity, and secretory functions in response to tumor challenge. Furthermore, by moving beyond binary “ON/OFF” dynamics, this titratable regulation may offer a distinct clinical advantage. It allows for the optimization of antitumor robust responses while minimizing the risk of toxicity.

### CSN CAR-T cells are potentially more persistent and less exhausted compared with conventional CAR-T cells *in vitro* culture

In CAR-T cells, CAR expression level typically declines over time, even without tumor stimulation, compromising both the efficacy and durability of therapy. To test whether our designed sCAR system could delay this loss, we cultured CAR-T cells and monitored CAR^+^ percentage and expression level (MFI) (Figures 5A and 5B). Each week, CAR-T cells were analyzed for CAR expression. Conv. CAR displayed a marked reduction in CAR^+^ percentage and levels, consistent with previous reports. By week 3, the CAR-positive fraction dropped around 40%, while expression level fell approximately two-thirds. In contrast, CSN-ON CAR-T cells maintained a more stable CAR^+^ percentage and delayed the decrease in CAR expression level. A stable CAR-positive population is a good indicator of persistence. The sCAR system delayed the decline in CAR expression during prolonged *in vitro* culture, suggesting the potential for improved functional maintenance compared with conventional CAR-T cells. To further compare persistence, we next assessed T cell exhaustion after tumor killing (Figures 5C and 5D). CSN-OFF CAR-T cells showed similarly low levels of exhaustion markers to NT, indicating the sCAR system may effectively prevent exhaustion in the OFF state. Importantly, upon tumor antigen stimulation, CSN-ON CAR-T cells exhibited markedly reduced expression levels of PD-1, TIM-3, and LAG-3 compared with Conv. CAR cells, under the tested *in vitro* stimulation conditions. These results suggest the potential for our switchable system may mitigate exhaustion while maintaining function (Figure 5C). T-SNE also visualized the distribution of exhaustion marker-expressing cells after tumor killing, providing a deeper understanding of T cell exhaustion-associated markers on the CAR-T cells (Figure 5D). Compared with Conv. CAR (left) and CSN-ON CAR (right) T cells, CSN-ON showed fewer exhausted subsets and lower levels of these exhaustion markers. We next performed an *in vitro* chronic stimulation model to evaluate whether CSN CAR-T cells preserve cytotoxic capacity after repeated stimulations, serving as a functional measure of potential persistence. CAR-T cells were challenged with Raji tumor cells for three rounds. After two rounds, CSN CAR-T cells were temporarily switched OFF for “rest”, then reactivated for a third stimulation (Figure 5E). We identified the tumor cytotoxicity after each round. CSN-ON CAR-T cells demonstrated robust cytotoxicity across all rounds, comparable to Conv. CAR, while also providing control in tumor killing (Figure 5F).

**Figure 5.**
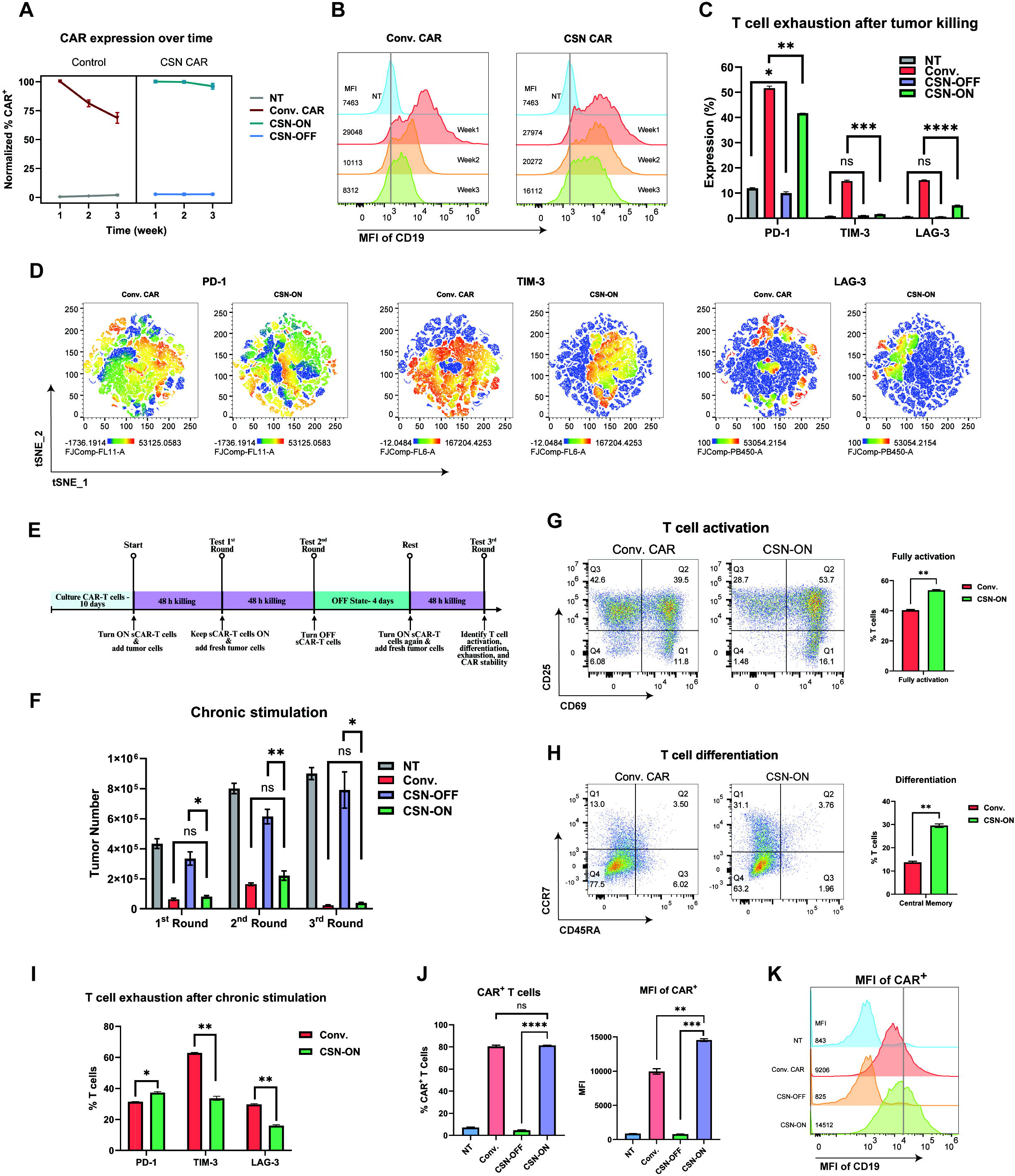
Characterization of persistence and exhaustion of CAR-T cells. (**A**) CAR expression rate over time. NT and Conv. were cultured with DMSO, while CSN CAR-T cells were incubated with either 5 μM GZR (ON state) or DMSO (OFF state). Functional CARs were detected weekly by staining with PE-CD19 protein and analyzed by flow cytometry. (**B**) CAR expression level (MFI) over time. Conv. were cultured with DMSO, and CSN CAR-T cells with 5 μM GZR (ON state). Expression level was measured as the median fluorescence intensity (MFI) of bound PE-CD19. (**C**) Exhaustion of CAR-T cells after tumor killing. CAR-T cells or NT cells were cocultured with Raji cells at a 1:1 E:T ratio for 4 days, then stained with antibodies against CD3, PD-1, TIM-3, and LAG-3 to detect exhaustion marker expression. (**D**) Flow cytometry analysis of exhaustion markers visualized by t-SNE. Marker expressions were gated on CD3^+^ T cells. PD-1 (CD3^+^PD1^+^); TIM-3 (CD3^+^TIM-3^+^); LAG-3 (CD3^+^LAG-3^+^). The color axis represents marker expression levels and the distribution of marker-positive cells. t-SNE-1 and t-SNE-2 represent the first and second dimensions of the t-SNE embedding and do not correspond to defined biological variables. (**E**) Timeline of chronic stimulation of CAR-T cells. CAR-T cells were cocultured with Raji cells for 48 h per round over three sequential rounds, following 10 days of manufacturing. NT, Conv., and CSN-OFF CAR-T cells were incubated with DMSO, while CSN CAR-T cells were turned ON with 5 μM GZR. E:T ratio was maintained at 1:1 for each round. T cell cytotoxicity was assessed after each round. (**F**) Cytotoxicity of CAR-T cells in the chronic stimulation model. Remaining tumor cells were quantified by flow cytometry and an automated cell counter. (**G**) Activation state of T cells after chronic stimulation. Cells were stained with CD3, CD25, and CD69 antibodies to show the activation state and level. Full activation was defined by the frequency of double-positive CD25^+^CD69^+^ T cells (Q2). (**H**) Differentiation of CAR-T cells after chronic stimulation. T cell subsets were analyzed by flow cytometry using CD3, CCR7, and CD45RA antibodies. Central memory T cells were defined as CD3^+^CCR7^+^CD45RA^−^ (Q1). (**I**) T cell exhaustion level after chronic stimulation. Expression levels of PD-1, TIM-3, and LAG-3 were measured by flow cytometry. (**J and K**) The percentage of CAR-positive T cells and CAR expression level after chronic stimulation. The positive CAR percentage and its expression level were measured by incubating cells with PE-CD19 protein in flow cytometry. Functional CARs bind with the fluorescent CD19 protein.

To gain deeper insight into the predictive factors of persistence, we profiled CAR-T cell activation, differentiation, exhaustion, and CAR expression following *in vitro* chronic stimulation. CSN-ON CAR-T cells displayed a higher proportion of CD69^+^CD25^+^ T cells (fully activated) than Conv. CAR, suggesting more robust yet controlled *in vitro* activation (Figure 5G). We next examined T cell differentiation, as central memory T cells are highly desired after tumor killing due to their longevity and proliferative capacity. CSN-ON CAR-T cells also showed a significantly higher proportion of central memory T cells than Conv. CAR, indicating that the switchable system promotes a more persistent and therapeutically favorable phenotype in the chronic stimulation model (Figure 5H). Because central memory T cells are less susceptible to exhaustion, we directly evaluated exhaustion-associated marker expression following chronic stimulation (Figure 5I). CSN-ON CAR-T cells exhibited markedly lower TIM-3 and LAG-3 levels than Conv. CAR, while PD-1 levels were slightly higher. Given that PD-1 is also a marker of antigen engagement, rather than terminal exhaustion markers like TIM-3 and LAG-3. These profiles suggested that CSN-ON CAR-T cells exhibited reduced expression of exhaustion-associated markers, including TIM-3 and LAG-3, under the tested *in vitro* stimulation conditions. Finally, we evaluated the stability of CAR expression after chronic stimulation (Figures 5J and 5K). CSN-ON CAR-T cells maintained a similar percentage of CAR-positive cells to Conv. CAR, but importantly, CSN CAR expression level remained significantly higher. In summary, CSN CAR-T cells demonstrated could facilitate stable CAR expression, reduced exhaustion-associated marker expression, sustained cytotoxicity across multiple *in vitro* tumor challenges, enhanced central memory formation, and potentially greater resistance to exhaustion compared to Conv. CAR-T cells under the tested model and conditions.

## Discussion

In this study, we developed an optimized, systematically characterized sCAR platform that enables reversible, graded, drug-controllable regulation of CAR expression, T-cell activation, and functions. By integrating the NS3/4A protease and its clinically approved inhibitors into the sCAR design, we established an optimized system that not only provides binary ON/OFF control but also enables dose-dependent tuning of CAR-T activity tested *in vitro*. Furthermore, screening three generations of NS3/4A inhibitors allowed us to identify small-molecule modulators with high efficacy, specificity, and low toxicity for this switchable CAR-T architecture. Collectively, our findings demonstrated that CSN CAR-T cells maintain potent antitumor activity, exhibit reduced exhaustion-associated marker expression, display potentially enhanced persistence, and may offer a safer therapeutic profile compared with conventional CAR-T. CSN CAR-T cells demonstrated lower expression of exhaustion-associated markers and higher central-memory frequencies after *in vitro* chronic stimulation. Interestingly, CSN-ON CAR-T cells occasionally exhibited activation and cytotoxicity comparable to or greater than conventional CAR-T cells despite sharing identical intracellular signaling domains. Although the precise mechanism remains unclear, the primary mechanism of regulation in the CSN platform is likely mediated through protease-dependent control of CAR surface availability. This effect may reflect differences in CAR surface expression dynamics, receptor turnover, or reduced tonic signaling prior to activation. Consistent with this interpretation, incorporation of the NS3/4A switch did not measurably alter basal CAR-T cell expansion. Furthermore, several factors may contribute to this effect. First, the ability to temporarily switch CAR-T cells OFF may reduce chronic antigen stimulation, which is a known driver of T cell exhaustion(38). In the OFF state, sCAR-T cells are shielded from excessive signaling, allowing recovery from activation-induced stress, and transient rest of CAR-T cells by switching it OFF may also restore CAR functions(34). Second, the graded control enabled by our system may allow for tuning of activation intensity, thereby avoiding overstimulation that can drive terminal exhaustion. Consistent with this, repeated tumor rechallenge assays revealed that CSN CAR-T cells maintained cytotoxicity, sustained CAR expression, and preserved memory-like features over multiple rounds of antigen exposure. While these observations may suggest a persistence-supportive phenotype, definitive evidence for improved durability requires longitudinal functional assays and *in vivo* validation. Moreover, CSN CAR-T cells may mitigate T-cell-associated toxicities by enabling precise regulation of CAR expression, T cell activation, cytokine release, and cytotoxicity, avoiding overactivation and delaying T-cell exhaustion. Additionally, in PBMC co-cultures, normal B cells were spared from cytotoxicity only when the sCAR system was switched OFF, confirming that pharmacologic suppression of surface CAR expression could abrogate on-target, off-tumor activity and prevent unintended elimination of healthy B cells in a mixed hematopoietic environment after tumor clearance. The goal is not to switch CAR-T cells OFF whenever remission is reached, but to offer a way to individualize the depth and duration of B-cell aplasia in selected patients. For the mechanism of surface CAR downregulation after turning OFF, the kinetics of CAR reversibility likely reflect a combination of intracellular inhibitor clearance, restoration of NS3/4A protease activity, and natural turnover of surface CAR molecules. The observed differences in downregulation rates among ASV, GZR, and GLE incubated CSN CAR-T cells *in vitro* further suggest that inhibitor-specific dissociation kinetics may contribute to the timing of protease reactivation following washout. Together, these processes govern the dynamic balance between CAR synthesis, intracellular cleavage, and surface CAR turnover, thereby enabling reversible control of receptor display.

Some traditional CAR-T therapies, such as tisagenlecleucel(4) and axicabtagene ciloleucel(2) are limited by uncontrolled activity, therapy-induced toxicities such as CRS, and eventual loss of persistence(11). To address these limitations, several switchable CARs have been reported. Most rely on adapter molecules that link tumor antigens to CAR scaffolds(39-41) or on small-molecule controlled dimerization of signaling domains(42). While these approaches afford temporal control and typically retain ligand-binding domains at the cell surface or modulate intracellular signaling while leaving surface CARs intact, which may permit tumor antigen engagement, trigger tonic signaling during OFF periods, or mask tumor antigens, blocking following immune recognition. In contrast, our strategy fundamentally differs from previously reported switchable CARs, as it directly regulates surface CAR receptor expression, preventing CAR binding in the OFF state. This provides tighter control and minimizes unintended antigen engagement. An additional consideration for protease-regulated CAR systems is whether cleaved CAR fragments interfere with tumor antigen recognition. Based on the known intracellular processing activity of HCV NS3/4A protease(43), we believe cleavage in the CSN platform occurs predominantly during CAR maturation and trafficking prior to stable surface localization, thereby limiting extracellular accumulation of soluble scFv-containing fragments. Consistent with this interpretation, we observed no detectable masking of CD19 epitopes on Raji tumor cells cocultured with CSN-OFF CAR-T cells, as assessed with an anti-CD19 antibody that has substantial epitope overlap with the CAR scFv binding region. These findings suggest that the OFF-state CSN system potentially minimizes persistent antigen engagement and avoids detectable tumor antigen masking *in vitro* coculture. Coupled with inhibitor dose, this sCAR system provides a single-component, pharmacologically addressable control over both antigen binding and signaling strength. Safety remains a critical challenge for CAR-T therapy, with CRS among the most concerning complications. In our system, cytokine production, including TNF-α, IFN-γ, and IL-6, was tightly controlled *in vitro*. Importantly, because clinical CRS is driven primarily by host myeloid cell-derived inflammatory signals (such as IL-6), we further evaluated macrophage-associated cytokine responses (IL-6) using a triple-culture system containing CAR-T cells, tumor cells, and primary human macrophages. In OFF and low-dose ON states, CSN groups showed substantially lower IL-6 secretion levels than conventional CAR-T cells, supporting the concept that graded pharmacologic regulation of CAR activity can attenuate macrophage-associated IL-6 *in vitro*. While these findings do not fully recapitulate the complexity of clinical CRS, they provide preliminary evidence that tuning CAR activation strength may reduce inflammatory signals associated with excessive immune activation. CSN CAR-T cells effectively prevented unintended cytokine release and fine-tuned cytokine secretion rather than abrupt overproduction *in vitro* culture conditions. Importantly, IL-6, a key mediator of CRS, plateaued at low drug concentrations, may enable a safety advantage by reducing the risk of CRS. The reversible and titratable nature of the CSN platform may enable proactive modulation of CAR-T activity to reduce the likelihood of excessive cytokine release and prolonged overactivation. The system may be particularly valuable for prophylactic dose-tuning and sustained management of CAR-T functional intensity.

We interpret these data as evidence that CSN permits dose-range-limited tuning of inflammatory outputs *in vitro*. Beyond cytokine management, the reversibility of our system may permit prompt attenuation or restoration of activity as clinically needed and has the potential to provide an operational safety lever compared with permanently active CAR-T cells. Importantly, GZR, one of the primary inhibitors used in this study, is an FDA-approved antiviral agent for hepatitis C virus infection with well-characterized clinical pharmacokinetics and safety profiles. This regulatory background provides translational support for its use as a pharmacological control element in the CSN CAR-T platform. In our study, short-term exposure to NS3/4A inhibitors did not induce detectable cytotoxicity in CAR-T cells, however, the effects of prolonged or repeated exposure on long-term T-cell expansion and persistence warrant further investigation. Our findings have several implications for the future of CAR-T therapy. First, the tunable CSN CAR-T cell system could enable more personalized dosing strategies, where the intensity of CAR-T activity is tailored to individual patient needs. For example, low drug concentrations could be used during initial treatment to reduce toxicities, followed by escalation to achieve deeper tumor clearance. In addition, administration of GZR in NSG mice successfully induced CAR surface expression *in vivo*, supporting the feasibility of pharmacologic regulation under physiologically relevant conditions. As for the dosage administration, because the percentage of induced CAR-positive cells directly determines the effective effector population, therapeutic activity may be achievable without requiring the highest dosage and maximal CAR induction. Nevertheless, detailed PK studies will be necessary to define clinically optimal dosing strategies and exposure windows for future translational applications. Secondly, reversible ON/OFF control provides a built-in safety mechanism, its reversible ON/OFF control may enable rapid attenuation or temporary suspension of activity if such effects arise during solid-tumor targeting, potentially limiting severity rather than preventing occurrence. Third, the predictive persistence and memory-like features may extend the durability of remission, addressing one of the major unmet needs in CAR-T therapy today. In summary, this design raises the possibility of developing next-generation CAR-T cells, may address key limitations, and represents promising steps toward safer and more durable cell-based immunotherapies. Despite these promising findings, several limitations of our study should be acknowledged. While they provide strong mechanistic insights, *in vivo* studies are needed to confirm persistence and efficacy in more physiologically relevant settings. An additional consideration for clinical translation is the potential immunogenicity of the HCV-derived NS3/4A module. Because NS3/4A is a viral protein, host immune recognition could potentially contribute to reduced persistence or immune-mediated clearance of engineered CAR-T cells in immunocompetent settings. In the current study, we demonstrated that administration of NS3/4A inhibitors successfully induced CAR expression *in vivo*, supporting the feasibility of drug-mediated regulation under physiological conditions. However, these studies were performed in immunodeficient mice and therefore do not evaluate adaptive immune responses against the NS3/4A module. Future studies in more immunologically relevant models will be important to determine the long-term immunogenicity of the CSN CAR system *in vivo*. Additionally, our work focused on CD19-targeting CAR, the system performance with other antigens or in solid tumor contexts remains to be determined. Therefore, future work should extend these findings into preclinical animal models to validate the persistence, exhaustion-resistance, and safety benefits of CSN CAR-T cells *in vivo*. Expanding this system to target solid tumors could further enhance its therapeutic potential. Altogether, this design raises the possibility of building next-generation CAR-T therapies with unprecedented control.

In conclusion, we presented a drug-addressable, reversibly switchable CAR-T platform that enables precise, drug-dependent control of surface CAR receptor expression and functions tested *in vitro*. Compared with Conv. CAR-T cells, our CSN CAR-T cells suggest the potential to exhibit reduced exhaustion, may enable enhanced persistence, and improved safety. By offering both reversibility and graded modulation, this system addresses key limitations of current CAR-T therapies and represents a promising step toward safer and more durable cell-based immunotherapies.

## Declarations

### Ethics approval and consent to participate

All animal experimental protocols were reviewed and approved by the Institutional Animal Care and Use Committee (IACUC) of Texas A&M University (Protocol No. 2025-0218 H). Primary human peripheral blood mononuclear cells (PBMCs) used in this study were purchased from a commercial vendor (STEMCELL Technologies, Vancouver, BC, Canada).

### Consent for publication

Not applicable.

### Availability of data and materials

Original data are available on request from the corresponding author, Wenshe Ray Liu.

### Authorship Contributions

W.R.L., Z.A.Z. and W.C. designed research; Z.A.Z. performed major parts of experiments, L.H., and T.H.S. performed experiments; Z.A.Z., Y.H., and X.S. analyzed data; Z.A.Z. and W.R.L. wrote the original draft, Z.A.Z., W.R.L., and W.C. revised the manuscript.

## Acknowledgements

We thank Dr. Andras Attila Heczey, a physician-scientist and full-time faculty member in the Department of Pediatrics of Baylor College of Medicine, for generously sharing the retro-vector backbone. We thank Dr. Peter J. A. Davies (faculty at Texas A&M University), Dr. Jason T. George (faculty at Texas A&M University), and Dr. Zhi Zachery Geng (postdoc at Texas A&M University) for providing training and helping with project design. The authors acknowledge TAMU-IBT flow cytometry and cell sorting core, directed by Dr. Margie Moczygemba.

