## Supplemental Tables and Figures for "Precise Control of Switchable Chimeric Antigen Receptor T Cells Allows Enhanced Safety and Less T Cell Exhaustion"

Supplementary Information for

### Precise Control of Switchable Chimeric Antigen Receptor T Cells Enables Graded Regulation of T-Cell Activity and Attenuates Exhaustion Markers

**Authors:** Zihan Anna Zhang[1], Logan Herring[1], Thuzar Hla Shwe[2], Yue Hu[1], Xiaotong Song[1], Wenyue Cao[2,*], and Wenshe Ray Liu[1,2,3,4,5,*]

[1]Institute of Biosciences and Technology and Department of Translational Medical Sciences, College of Medicine, Texas A&M University, Houston, TX 77030, USA

[2]Texas A&M Drug Discovery Center and Department of Chemistry, College of Arts and Sciences, Texas A&M University, College Station, TX 77843, USA

[3]Department of Biochemistry and Biophysics, College of Agriculture and Life Sciences, Texas A&M University, College Station, TX 77843, USA

[4]Department of Cell Biology and Genetics, School of Medicine, Texas A&M University, College Station, TX 77843, USA

[5]Department of Pharmaceutical Sciences, Irma Lerma Rangel College of Pharmacy, Texas A&M University, College Station, TX 77843, USA

*To whom correspondence may be addressed: Wenyue Cao and Wenshe Ray Liu

Table of Contents

Supplementary Tables … P3-P5

Supplementary Figures … P6-P11

**Table S1: Reagent and Resource**

| REAGENT or RESOURCE | SOURCE | Identifier |
|---|---|---|
| **Antibodies** | | |
| PE anti-human CD3 | BioLegend | 300441 |
| Pacific Blue anti-human CD223 | BioLegend | 369341 |
| FITC anti-human CD366 (Tim-3) | BioLegend | 345021 |
| APC anti-human CD279 (PD-1) | BioLegend | 379207 |
| APC/Cyanine7 anti-human CD3 | Biolegend | 300426 |
| APC anti-human CD69 | Biolegend | 310909 |
| APC/Cyanine7 anti-human CD25 | Biolegend | 302613 |
| Brilliant Violet 510 anti-human IL-2 | Biolegend | 500338 |
| APC anti-human/mouse Granzyme B Recombinant | Biolegend | 372203 |
| APC/Cyanine7 anti-human Perforin | Biolegend | 308127 |
| APC anti-human CD197 (CCR7) | Biolegend | 353213 |
| Brilliant Violet 570 anti-human CD45RA | Biolegend | 304131 |
| FITC Mouse anti-human CD3 | BD Biosciences | 555332 |
| PE anti-human CD19 | Biolegend | 302208 |
| APC antihuman CD19 (SJ25C1) | Biolegend | 363005 |
| Brilliant Violet 421 anti-human CD56 (NCAM) | Biolegend | 318327 |
| PE/DazzleTM 594 anti-human CD14 | Biolegend | 325633 |
| F(ab')$_2$ Fragment Rabbit Anti-Mouse IgG (H+L) | JacksonImmunoResearch | 315-006-003 |
| **Chemicals and recombinant proteins** | | |
| Asunaprevir | MCE | HY-14434 |
| Danoprevir | MCE | HY-10238 |
| Simeprevir | MCE | HY-10241 |
| Tipranavir | MCE | HY-15148 |
| Vaniprevir | AdooQ Biocience | A11600 |
| Boceprevir | MCE | HY-10237 |
| Glecaprevir | MCE | HY-17634 |
| Narlaprevir | MCE | HY-10300 |
| Grazoprevir | MCE | HY-15298 |
| PE-CD19 protein | ACRO Biosystems | CD9-HP2H5 |
| Recombinant Human IL-2 | PeproTech | 200-02 |
| GolgiStop Protein Transport Inhibitor (Monensin) | BD Biosciences | 554724 |
| Helix NP Blue | Biolegend | 425305 |
| Dynabead Human T-Activator CD3/CD28 | ThermoFisher Scientific | 11131D |

| | | |
|---|---|---|
| Polyethylene Glycol 8000 (PEG) | ThermoFisher Scientific | BP233-100 |
| Polybrene | Millipore sigma | TR-1003-G |
| PEI-max | Polysciences | 49553-93-7 |
| M-CSF | PeproTech | 300-25-50UG |
| PEG-300 | MCE | HY-Y0873 |
| Tween 80 | MCE | HY-Y1891 |
| ACK buffer | Gibco | A1049201 |
| **Critical commercial assays** | | |
| ClinMax Human IL-6 ELISA Kit | ACRO Biosystems | CRS-B001 |
| ELISA MAX set- Human TNF-α | Biolegend | 430204 |
| ELISA MAX set- Human IFN-γ | Biolegend | 430104 |
| **Experimental models: Cell lines** | | |
| HEK 293T cells | ATCC | CRL-3216 |
| K562 cells | ATCC | CCL-243 |
| Raji cells | ATCC | CCL-86 |
| Human Peripheral Blood Mononuclear Cells | STEMCELL | 70025.2 |
| **Recombinant DNA** | | |
| CSC CAR | This paper | N/A |
| CSN CAR | This paper | N/A |
| C&N CAR | This paper | N/A |
| **Software and algorithms** | | |
| FlowJo 10.10.0 | FlowJo | https://www.flowjo.com/ |
| GraphPad Prism 10.6.1 | GraphPad | https://www.graphpad.com |
| BioRender | Biorender | https://www.biorender.com/ |
| Adobe Illustrator 29.8 | Adobe | https://www.adobe.com/ |

**Table S2: Sequence Information**

| **Gene fragment** | **DNA Sequence** |
|---|---|
| CAR receptor (scFv, anti-hCD19) | ATTCCAGACATTCAGATGACGCAAACAACGTCAAGCCTCTCTGCATC<br>ACTGGGTGATAGGGTAACGATAAGTTGTAGAGCGTCACAGGATATCT<br>CCAAATACTTGAACTGGTACCAGCAGAAGCCAGACGGGACTGTAAA<br>ACTCCTCATCTACCATACTTCTCGACTCCACTCCGGTGTGCCCAGTAG<br>ATTTTCAGGATCCGGCAGTGGGACAGATTATTCTCTTACAATTAGCA<br>ACTTGGAACAGGAGGACATCGCAACTTACTTCTGCCAGCAGGGAAAC<br>ACGCTCCCTTATACGTTCGGGGGAGGGACGAAACTCGAAATCACT |
| NS3/4A switch | ACTGGATGTGTTGTGATAGTCGGTCGCATAGTACTCTCTGGCTCTGGA<br>ACATCTGCCCCAATTACGGCGTATGCGCAGCAGACACGAGGACTTCT<br>GGGTTGCATTATCACGAGCCTGACGGGGAGAGACAAGAATCAGGTG<br>GAGGGGGAGGTACAGATAGTAAGCACGGCAACGCAAACGTTCCTTG<br>CGACATGCATAAATGGAGTTTGCTGGGCTGTCTACCACGGAGCAGGA<br>ACCCGCACAATTGCGTCACCTAAAGGCCCGGTAATACAGATGTACAC<br>AAACGTTGACCAGGACCTTGTTGGGTGGCCCGCGCCGCAAGGGAGTC<br>GATCCCTGACTCCGTGTACGTGCGGAAGCTCTGATTTGTATCTTGTGA<br>CACGCCACGCGGACGTGATTCCCGTCCGACGCCGAGGGGATTCTCGC<br>GGAAGTCTGTTGAGCCCACGGCCAATATCTTACCTTAAAGGTAGTAG<br>CGGGGGCCCTCTGTTGTGTCCTGCGGGGCACGCCGTGGGTCTTTTTCG<br>GGCAGCAGTCTGTACTAGGGGTGTAGCCAAGGCCGTGGACTTTATTC<br>CTGTAGAAAACCTCGAGACTACCATGCGGTCCCCGGTGTTCACAGAT<br>AATTCATCTCCTCCCGCTGTCACTCTCACCCAT |
| CSC/CSN | GACGAAATGGAGGAATGTTCTCAGCAC |
| TM | TTTTGGGTTTTGGTAGTCGTGGGCGGCGTGCTGGCATGCTATTCACTG<br>CTGGTAACCGTTGCATTCATCATCTTCTGGGTG |
| CD28 | AGATCAAAACGGTCAAGACTGCTCCATAGTGATTATATGAATATGAC<br>GCCAAGGAGACCTGGTCCAACCCGGAAGCATTATCAACCCTACGCAC<br>CACCTAGGGATTTCGCTGCTTATAGGTCA |
| 4-1BB | AAGAGAGGAAGGAAGAAGTTGCTCTACATTTTTAAACAGCCCTTCAT<br>GAGGCCTGTACAGACAACCCAGGAGGAGGACGGGTGTAGTTGTAGA<br>TTTCCGGAGGAAGAGGAAGGCGGCTGTGAGCTC |
| CD3ζ | CGCGTAAAATTTAGCAGGTCTGCGGATGCCCCAGCGTACCAGCAAGG<br>CCAGAATCAGTTGTACAATGAGCTTAACCTTGGAAGACGCGAGGAGT<br>ACGACGTACTCGATAAGAGGAGGGGAAGGGATCCAGAGATGGGTGG<br>CAAACCACGCCGAAAAAACCCTCAGGAAGGACTGTATAACGAACTC<br>CAGAAAGATAAGATGGCTGAAGCTTATAGTGAGATAGGAATGAAGG<br>GAGAAAGACGCCGAGGAAAGGGACATGACGGTCTTTACCAGGGCTT<br>GAGCACTGCGACTAAAGACACCTACGATGCGCTCCATATGCAGGCTC<br>TGCCTCCGAGA |

**Supplementary Figures**

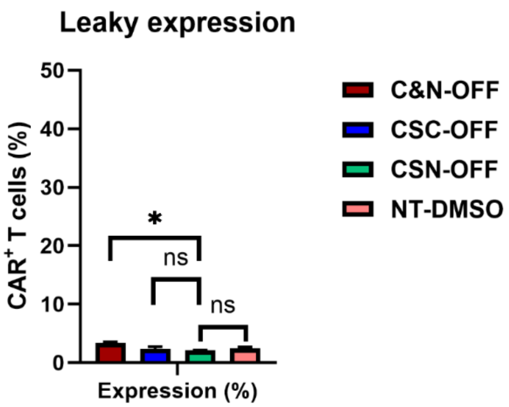


**Figure S1. Comparison of basal leakage among the C&N, CSC, and CSN CAR designs.** Basal sCAR surface expression in the OFF state was evaluated by flow cytometry. C&N, CSC, and CSN CAR-T cells were cultured with DMSO for 24 h to maintain the OFF condition. Among the three designs, CSN and CSC CARs exhibited lower basal CAR expression, which showed minimal leakage comparable to NT cells. The comparison between CSN-OFF and NT-DMSO groups was not statistically significant ($p = 0.257$).

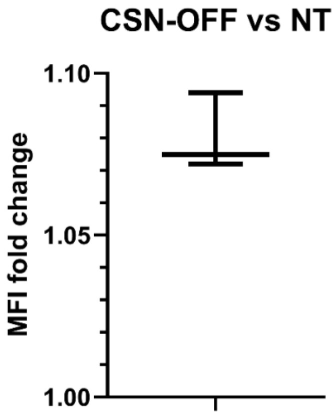


**Figure S2. MFI fold-change comparison between CSN-OFF and NT cells.** CSN CAR-T cells and NT T cells were cultured with DMSO for 24 h under the OFF condition. Surface CAR intensity was analyzed by flow cytometry, and mean fluorescence intensity (MFI) fold change was calculated relative to NT cells. The mean MFI fold change of CSN-OFF cells was approximately 1.08 compared with NT cells, indicating minimal basal signal above background levels.

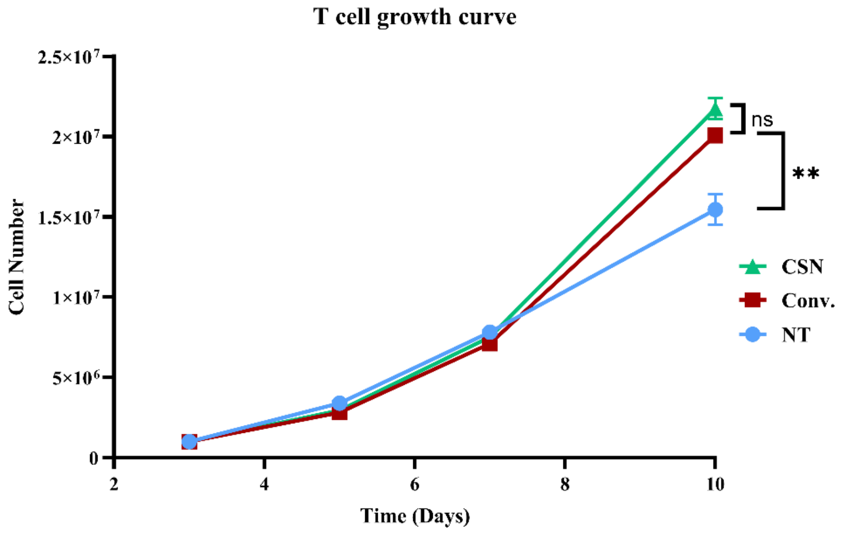


**Figure S3. In vitro cell growth of T cells and CAR-T cell**s. T cells, Conv. CAR-T cells and CSN CAR-T cells were cultured *in vitro*, and total cell numbers were measured every other day using an automated cell counter. CSN CAR-T cells exhibited a growth rate comparable to Conv. CAR-T cells, while both CAR-T groups expanded more rapidly than NT T cells.

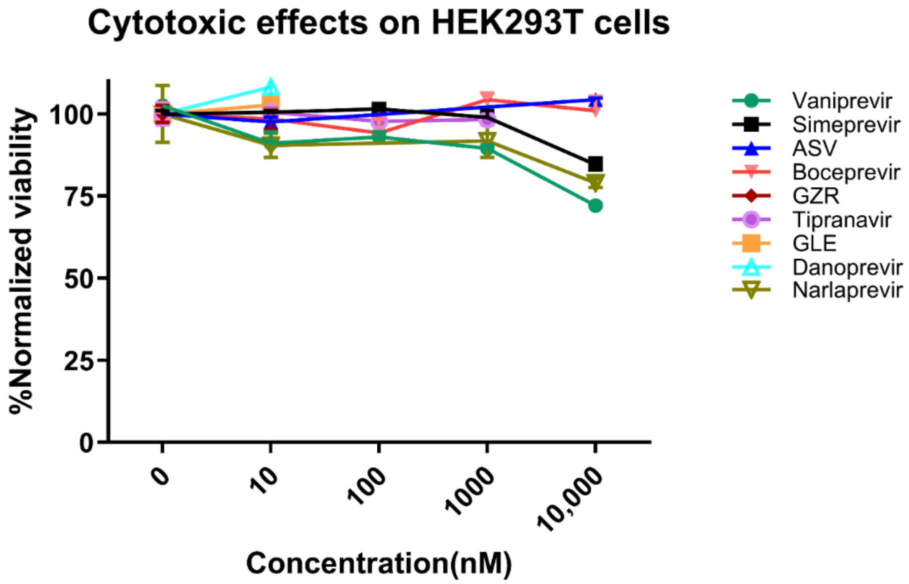


**Figure S4. The toxicity of the nine small-molecule drugs on HEK293T cells.** Toxicity was measured using MTT assays. HEK293T cells were incubated with varying concentrations of the nine small molecules for 48 h. After incubation, the cell viability was measured by MTT.

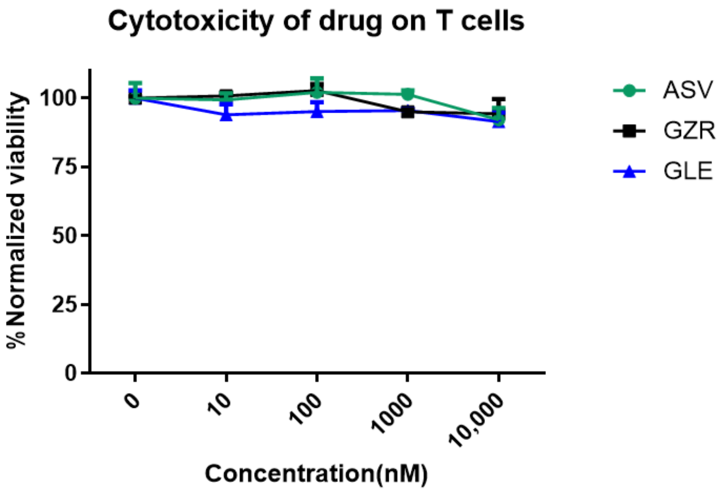


**Figure S5. The toxicity of the "top 3" small-molecule drugs on human T cells.** Toxicity was measured using MTT assays. Human T cells were incubated with varying concentrations of the three small molecules for 48 h. After incubation, the cell viability was measured by MTT.

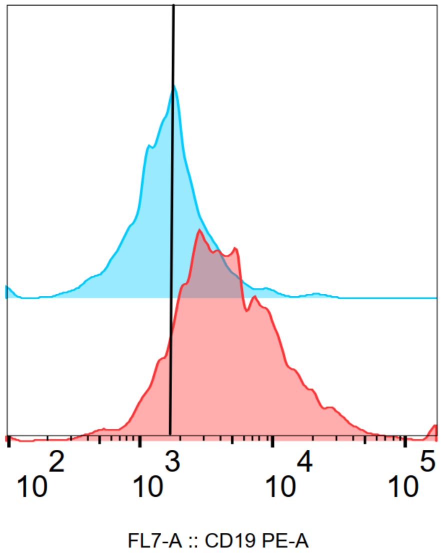


**Figure S6. Drug-induced CAR surface expression *in vivo* in NSG mice.** CSN CAR-T cells were intravenously infused into NSG mice at 5×10^6 CAR+ T cells per mouse. Peripheral blood was collected 2 days after infusion and analyzed for baseline CAR expression (blue). Mice (n=3) then received grazoprevir (GZR) treatment at 50 mg/kg by daily intraperitoneal injection for 2 consecutive days, followed by repeat peripheral blood collection for CAR expression analysis (red). Mouse PBMCs were stained with anti-human CD3 antibody and PE-conjugated CD19 protein to detect CAR expression on human T cells. CAR expression levels were quantified as MFI within the hCD3+ populations.

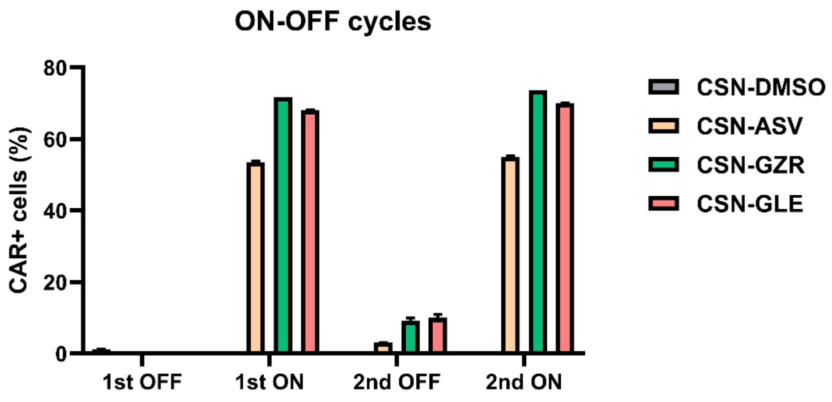


**Figure S7. Reversible ON/OFF cycling of CAR expression in CSN CAR-T cells.** CSN CAR-T cells were initially cultured without controller drugs to maintain the OFF state (CSN-DMSO). To induce the ON state, cells were treated with ASV, GZR, or GLE for 48 h (1st ON). Cells were then washed and cultured in drug-free medium for 72 h to return to the OFF state (2nd OFF). Subsequently, the same inhibitors were re-administered for an additional 48 h to induce a second ON state (2nd ON). Surface CAR expression was analyzed by flow cytometry using PE-conjugated CD19 protein staining, and CAR-positive populations were quantified.

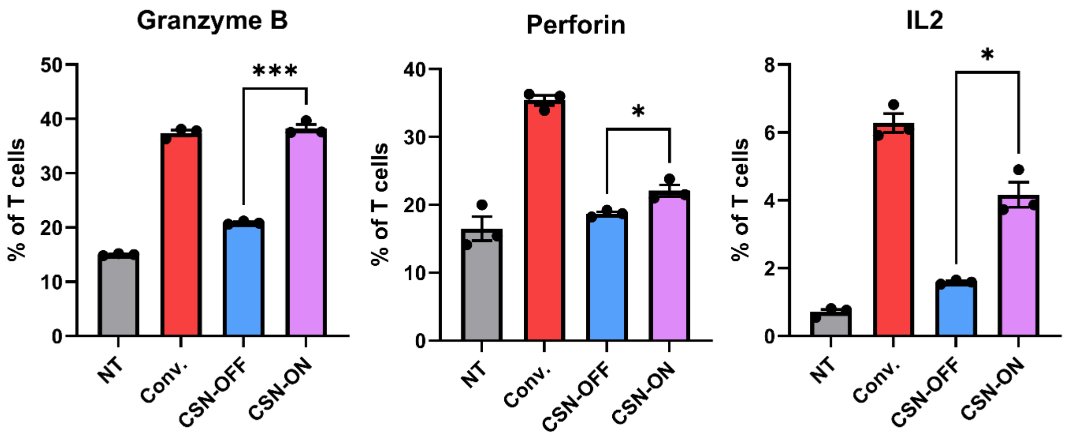


**Figure S8. The control of cytotoxic effector molecule expression on CSN CAR- cells.** All group cells were incubated with Raji tumor cells at a 1:1 E:T ratio. NT, Conv., and CSN-OFF CAR-T cells were incubated with DMSO, and CSN-ON cells were incubated with 5 μM GZR. After coculturing with Raji and DMSO or GZR, the protein transport inhibitor was added to the media 4 h before analysis to block protein secretion and accumulate secretive proteins at the endoplasmic reticulum (ER). The cells were labeled with the CD3 antibody to identify T cells, then fixed and permeabilized. After permeabilization, cells were labeled with antibodies against granzyme B, perforin, and IL-2. The expression of cytotoxic effector molecules was analyzed in flow cytometry.

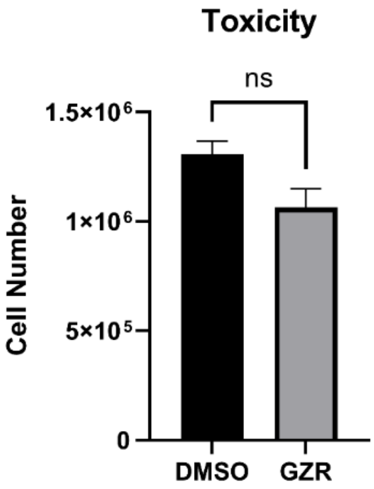


**Figure S9. Effect of grazoprevir on K562 tumor cell viability.** K562 cells were cultured in the presence or absence of 5 μM GZR for 48 h. Following incubation, total cell numbers were quantified using an automated cell counter to assess potential direct effects of GZR on tumor cell viability and growth.

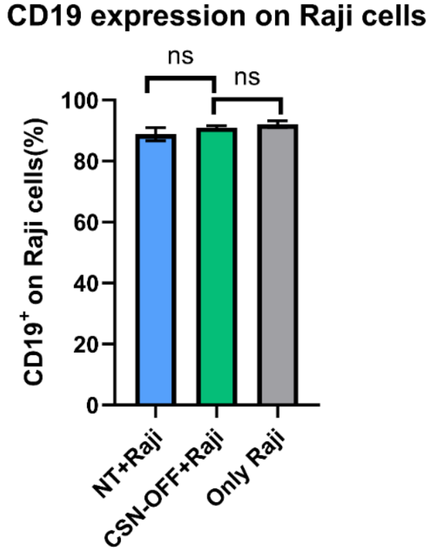


**Figure S10. Evaluation of tumor antigen masking by CSN-OFF CAR-T cells.** Raji tumor cells were cocultured with NT cells or CSN-OFF CAR-T cells for 24 h. Following coculture, CD19 epitope accessibility on Raji cells was evaluated by flow cytometry using an anti-CD19 antibody (SJ25C1 clone) with substantial epitope overlap with the CAR scFv recognition region. No detectable reduction in CD19 antibody binding was observed in the CSN-OFF group compared with NT or Raji-only controls, indicating minimal tumor antigen masking by CSN CAR-T cells in the OFF state.

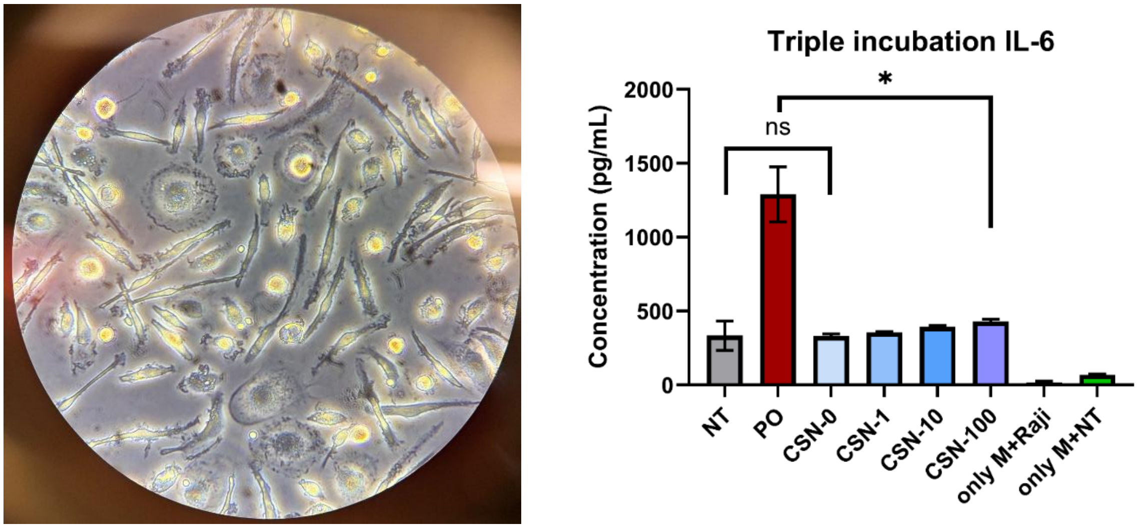


**Figure S11. Macrophage-associated IL-6 production in a triple culture CRS model.** Primary human macrophages (Left, bright field, 40x objective lens) were cocultured with Raji tumor cells and CAR-T cells at a 1:1:1 ratio under the indicated drug treatment conditions. The NT group was treated with DMSO, and the positive control (PO, conventional CAR-T cells) group was treated with GZR. For the CSN groups, CSN-0 was cultured with DMSO, while CSN-1, CSN-10, and CSN-100 were treated with 1 nM, 10 nM, and 100 nM GZR, respectively, for 24 h. Additional control groups included macrophages cocultured only with Raji cells (only M+Raji) or only with NT T cells treated with DMSO (only M+NT). Following coculture, supernatants were collected, and IL-6 concentrations were quantified by ELISA (Right) to evaluate macrophage-associated IL-6 levels under different levels of CSN CAR activation.
